# Real-time whole-plant dynamics of heavy metal transport in *Arabidopsis halleri* and *Arabidopsis thaliana* by gamma-ray imaging

**DOI:** 10.1101/428417

**Authors:** Kaisa Kajala, Katherine L. Walker, Gregory S. Mitchell, Ute Krämer, Simon R. Cherry, Siobhan M. Brady

**Affiliations:** Department of Plant Biology and Genome Center, University of California Davis, Davis CA, USA; Plant Ecophysiology, Institute of Environmental Biology, Utrecht University, Utrecht, the Netherlands; Department of Biomedical Engineering, University of California Davis, Davis CA, USA; Molecular Genetics and Physiology of Plants, Ruhr University Bochum, Bochum, Germany

**Keywords:** *Arabidopsis halleri*, metal hyperaccumulation, metal transport, metal uptake, nuclear imaging, SPECT, zinc

## Abstract

Heavy metals such as zinc are essential for plant growth, but toxic at high concentrations. Despite our knowledge of the molecular mechanisms of heavy metal uptake by plants, experimentally addressing the real-time whole-plant dynamics of heavy metal uptake and partitioning has remained a challenge. To overcome this, we applied a high sensitivity gamma-ray imaging system to image uptake and transport of radioactive ^65^Zn in whole-plant assays of *Arabidopsis thaliana* and the Zn hyperaccumulator *A. halleri*. We show that our system can be used to quantitatively image and measure uptake and root-to-shoot translocation dynamics of zinc in real time. In the metal hyperaccumulator *Arabidopsis halleri*, ^65^Zn uptake and transport from its growth media to the shoot occurs rapidly and on time scales similar to those reported in rice. In transgenic *A. halleri* plants in which expression of the zinc transporter gene *HMA4* is suppressed by RNAi, ^65^Zn uptake is completely abolished.

**HIGHLIGHT:** We have used gamma-ray imaging to visualize the stark differences of real-time whole-plant dynamics of zinc root-to-shoot transport in heavy metal hyperaccumulating and non-accumulating *Arabidopsis*.

## INTRODUCTION

Most plants actively prevent the accumulation of high levels of metals in their aboveground biomass through a functional network of metal homeostasis, in order to avert toxicity. However, metal hyperaccumulators have developed rare adaptations to their environments and selectively extract specific metals from the soil and accumulate them in their shoots at very high concentrations without incurring symptoms of toxicity (Baker and Brooks, 1989; Frérot *et al*., 2010). Metal hyperaccumulator plant species accumulate one or several metallic or metalloid elements in aboveground biomass at concentrations above a threshold that is two to three orders of magnitude higher than in leaves of most species on normal soils, and at least one order of magnitude greater than the usual range found in plants from metalliferous soils, in their natural habitat (Pollard *et al*., 2014). Metal hyperaccumulation in plants can have commercial applications including phytostabilization (revegetation of contaminated soils), phytomining (extraction of metals from plants for their value) or phytoremediation (plant-based remediation of soils).

*Arabidopsis halleri* (formerly known as *Cardaminopsis halleri*) is a metal hyperaccumulator species and a facultative metallophyte, i.e., it grows naturally on both metal-contaminated toxic soils and pristine soils. That is, it is able to grow on both metalliferous as well as non-metalliferous soils. Zinc (Zn) hyperaccumulation (at least 10,000 μg of Zn per g of dry leaf tissue) in *A. halleri* is species-wide, whereas cadmium (Cd) hyperaccumulation (at least 300 μg of Cd per g of dry leaf tissue) is geographically confined (Bert *et al*., 2000, 2003; Pauwels *et al*., 2006; Frérot *et al*., 2010). The molecular mechanisms underlying metal hyperaccumulation in *A. halleri* are only partly understood. Typically Zn^2+^ and other poorly soluble transition metal cations are mobilized in the soil through acidification of the rhizosphere and the release of organic chelators by the root. These transition metal ions are then taken up into the root symplasm through ZIP transporters (ZRT, IRT-like Protein) located in the epidermal plasma membrane (Grotz *et al*., 1998; Ramesh *et al*., 2003). They are then thought to move symplastically from cell to cell and are eventually loaded into the apoplastic xylem through the transporters <u>H</u>EAVY <u>M</u>ETAL <u>A</u>TPASE 4 (HMA4) and HMA2 (Hussain *et al*. 2004). A quantitative trait locus for Cd^2+^ hypertolerance in *A. halleri* was found to map to a chromosomal region containing the *HMA4* gene (Courbot *et al*., 2007). Transgenic *A. halleri* lines in which *HMA4* was silenced by RNA interference were employed to demonstrate a key role for *HMA4* in Zn and Cd hyperaccumulation as well as Zn and Cd hypertolerance (Hanikenne *et al*., 2008). In comparison to *A. thaliana*, the strongly enhanced expression of *HMA4* in *A. halleri* results from a combination of modified *cis*-regulatory sequences and copy number expansion (Hanikenne *et al*., 2008). Transfer of an *A. halleri HMA4* gene into *A. thaliana* recapitulates Zn partitioning into xylem vessels and the constitutive transcriptional upregulation of Zn uptake system-encoding Zn deficiency-responsive genes including *ZIP4* and *IRT3* in roots. Other genes associated with metal hypertolerance and/or hyperaccumulation in *A. halleri* include constitutively highly expressed *<u>N</u>ICOTIAN<u>A</u>MINE <u>S</u>YNTHASE* (*NAS2* and *NAS3*), *<u>M</u>ETAL <u>T</u>OLERANCE <u>P</u>ROTEIN 1* (*MTP1*) and *<u>N</u>ATURAL <u>R</u>ESISTANCE-<u>A</u>SSOCIATED <u>M</u>ACROPHAGE <u>P</u>ROTEINS* (*NRAMP*) genes (Krämer *et al*., 2007; Hanikenne and Nouet, 2011).

The functional characterization of candidate metal transporters can reveal the molecular mechanisms by which metal hyperaccumulation occurs. Imaging methods can reveal the spatiotemporal dynamics of metal translocation within and between roots and shoots. According to the present model, hyperaccumulation is primarily a consequence of high rates of metal loading into the root xylem, which depletes the metal in the root symplasm and secondarily induces elevated rates of metal uptake into the root (Hanikenne *et al*., 2008; Krämer 2010). Additional contributions were proposed from chelation in the root symplasm (Deinlein *et al*., 2012), high rates of unloading from the xylem, and distribution across leaf tissues where the metal can be stored in mesophyll cells and is present in high amounts in vacuoles (Hanikenne *et al*., 2008). However, the time scale by which metal uptake into the root is followed by root-to-shoot translocation remains unknown. Within the root, mathematical modeling approaches have explored the importance of ZIP regulation, HMA abundance and symplastic transport in the creation of the radial pattern of Zn within primary roots of *A. thaliana* (Claus *et al*., 2013). Modeling predicted that Zn transport in the symplasm takes place at the same time scale as symplastic transport with Zn carried along the water flow path at the same velocity of water (Claus *et al*., 2013). Transpiration is thus a key determinant of the rate of this water transport for symplastic, root cell-to-cell transport. The metal must then be loaded into the xylem by the HMA4 transporter to undergo apoplastic transport from the root to shoot. In the hyperaccumulator *A. halleri*, Cd translocation from the external hydroponic medium into the xylem was very rapid: After a two-hour exposure to Cd, its concentration in xylem sap was 5-fold higher than that in the external solution (Ueno *et al*., 2008). This Cd concentration in xylem sap decreased with increasing the concentration of external Zn suggesting that it is a function of the HMA4 transporter. Based on nuclear magnetic resonance (NMR) and computational modeling of metal speciation by Geochem-PC of xylem sap, Cd was proposed to mainly occur in the free ionic form in the xylem sap.

Invasive methods involving decapitation of plants or detachment of stems have revealed aspects of the dynamics of apoplastic transport once metal is loaded into the xylem. Apoplastic transport of water in xylem of *A. thaliana* occurs within 30 min (Park *et al*., 2014), while xylem sap of *A. halleri*, collected after exposure to high Cd, accumulates Cd within the same time scale of 30 min (Ueno *et al*., 2008). At the single cell or tissue level, fluorescence resonant electron transfer (FRET) imaging in root cells revealed a high-affinity, low-capacity uptake system, a low-affinity, high-capacity uptake system as well as of a mechanism allowing Zn^2+^ release from internal cell stores in *A. thaliana* (Lanquar *et al*., 2014). Synchrotron-based techniques revealed that high concentrations of Cd were located in the vascular system of both the hyperaccumulator *A. halleri* and the non-accumulator, *A. lyrata* in the vascular system of the mid-rib and in secondary veins. The two species differ in the extent of their accumulation of Cd in the mesophyll, with *A. halleri* showing enriched levels relative to *A. lyrata* (Isaure *et al*., 2006).

To complement these often spatially narrow or invasive studies, whole plant imaging enables direct large-scale material transport studies, addressing spatiotemporal transport within the context of the entire intact plant, from root to shoot for a period of up to several days. Here, we use a high-sensitivity un-collimated detector single photon emission computed tomography (SPECT)-type imaging system (UCD-SPI (Walker *et al*., 2015)), to study Zn metal transport in a hyperaccumulator, *A. halleri*, and in a non-accumulator *A. thaliana*. We further examine the influence of the HMA4 transporter on Zn transport in *A. halleri* by use of an HMA4 knockdown line.

## MATERIALS AND METHODS

### Plant Material and Growth Conditions

All plants were cultivated in a growth chamber with a 16h:8h light:dark cycle at 22 °C and 50–75% humidity with a light intensity of 100 µmol m^−2^ s^−1^. We used *A. thaliana* (accession Columbia [Col-0]), *A. halleri* (LAN 5, Langelsheim, Germany), and *AhHMA4-*RNAi *A. halleri* (Line 4.2.1, Langelsheim, Germany) (Hanikenne *et al*., 2008). Seeds of *A. thaliana* Col-0 were grown to an age of 18 days on a mix of vermiculite and sand. 18-day-old plants were transferred from vermiculite and sand to a hydroponic Hoagland nutrient solution (Becher *et al*., 2004) without ZnSO_4_. Zn concentration in this medium was measured to be 0.12 µM (inductively coupled plasma-atomic emission spectrometry (ICP-AES), UC Davis Analytical Lab). The hydroponic growth was conducted in Magenta vessels (GA-7, Sigma Aldrich), each plant in an individual vessel, on a mesh square suspended over 150 ml of growth medium. Vegetatively growing *A. halleri* plants were replicated via cuttings and grown in hydroponics. The cuttings were first allowed to root on 1x SensiGrow (Advanced Nutrients, USA) nutrient solution for 14 days, before being transferred to Hoagland nutrient solution without Zn. The hydroponic solutions were changed every 4–5 days. Both *A. halleri* and *A. thaliana* plants were grown on Hoagland solution without Zn for 19–21 days prior to imaging to acclimate to Zn deprivation. Both *A. thaliana* and *A. halleri* were at mature vegetative growth stage during the imaging experiments.

### Imaging Protocol and Conditions

Zn isotope was obtained from the National Isotope Development Center (Oak Ridge National Laboratory), in the form of ZnCl^2^ in solution. The half-life of ^65^Zn is 243.9 days, and its primary gamma-ray decay product is of energy 1116 keV (51% branching ratio). A small positron emission branching ratio of 1.4% also exists.

Plants were transferred from the growth chamber to the imaging lab prior to imaging in batches of three where they grew during the time it took to image them. The imaging lab light level was 12 µmol m^−2^ s^−1^ PAR, with a light:dark cycle of 10h:14h, and was at temperature of 20 °C. Prior to imaging, each plant was transferred from the no-Zn Hoagland solution into a solution with radioactive Zn and 1 µM ZnSO_4_. The plants were incubated in a centrifuge tube containing 50 mL of Hoagland solution with 1 µM ZnSO_4_, to which a small amount of the ^65^Zn had been added. Each plant was held in a plastic funnel to avoid direct contact of the plant shoot with the radioactive solution, and incubated in the ^65^Zn solution for 60 min. The plant was placed in the funnel so that the roots were in the spout and the shoot was in the wide opening funnel mouth, and the funnel was placed in the centrifuge tube with the fluid level coming to the full level of the funnel spout to fully immerse the roots. To avoid generating a large amount of radioactive waste all plants were incubated in the same tube. For the first plant the ^65^Zn concentration was 7.2 nM and for the final plant the ^65^Zn concentration was 5.6 nM, corresponding to an activity range of 200 µCi to 150 µCi in the 50 mL tube. The concentration values for ^65^Zn in the incubation tube were determined by activity measurement of the tube in a calibrated well counter (Capintec, Inc., Florham Park, NJ). The uptake into each plant was typically 4 to 5 µCi.

Following incubation, each plant was rinsed through three washes of Hoagland solution (with 1 µM ZnSO_4_) to remove any (non-uptaken) radiolabelled media remaining on the root surface. This was performed with the plant still in the funnel and fresh 50 mL centrifuge tubes. Each rinse was of 20 seconds, during which the plant was gently swirled in the tube. The rinsing tubes were measured for activity rinsed off of the plant roots and only a small fraction was ever present. Final rinsing for each plant was always performed with a fresh solution. Rinsing solutions were one of the larger components of radiological waste which was generated. The imaging was carried out on Hoagland solution with 1 µM ZnSO_4_.

For gamma ray imaging on the UCD-SPI system, the plants were carefully removed from the funnel and gently constrained with a holder of two 10 cm × 15 cm pieces of thin transparent plastic, spaced to be separated by 2 cm. The narrow holder constrained the plant leaves to be close to the imaging system and to have the entire plant roots and shoot be within the system field of view. The holder contained the growth media at a level which completely covered the roots, a few cm below the top opening of the holder. Most of the leaves extended above the holder opening. Only one of the detector heads of the system was used in order to provide better monitoring of the plant condition throughout the imaging process. Plants were imaged continuously for a period of several hours until the gamma ray spatial distribution reached a steady state. Time periods of imaging ranged from 20 h to 70 h for the 11 plants.

Data were collected in files recording positions and energies of up to 8 × 10^6^ gamma ray events detected by the system. Given the efficiency and sensitivity of the system this corresponded to a time extent of 20 minutes for the larger plants with high Zn uptake, to multiple hours for the somewhat smaller *A. thaliana* plants. The data files were used as the time points for the subsequent analysis. For each plant, two non-overlapping regions of interest (ROIs) of area 1280 mm^2^ (16 by 20 pixels of detector, each of dimension 2 by 2 mm^2^) were defined on the detector area, corresponding to the roots and to the leaves of the plant. Total counts recorded in each area in a data file were normalized to the time duration of the data file, to result in a value for counts s^−1^ 1280 mm^−2^. Due to the lack of spatial resolution of the system, some counts in each ROI may have come from the opposite part of the plant, but given the close geometry and solid angle considerations, the fraction of the total is small. A total of eleven plants were imaged in the UCD-SPI system, three *A. thaliana*, and four each of the *A. halleri* genotypes.

### Quantification of Zn remaining in Growth Media after Imaging

At the end of plant imaging, two values were recorded from an electronic pulse counter unit (Tennelec TC512 Dual Counter Timer) for the overall trigger rate of the system: the final rate with the plant in place in the system, and the rate with the plant removed (but the growth media still remaining in the holder). Measurements of the plant components in the well counter were made post imaging but due to the low activity levels (often 2 µCi or lower) the measurements were not stable above background.

Parallel re-supply experiments with no radiolabel were done in triplicate to quantify the amount of Zn in the media before and after 24 h resupply of 1 µM Zn, and hence the ability of the plants to deplete Zn from their growth media. The Zn concentration in the growth media samples were quantified using ICP-AES analysis (UC Davis Analytical Lab). The re-supply experiments without radiolabel were carried out in Magenta boxes in the growth chamber for a 24 h period.

### Quantification of Transport from Imaging Data

Absolute sensitivity of the system is known from measurement with a centrifuge tube containing known ^65^Zn activity. The calibrated system performance as measured with point gamma ray sources of a variety of energies broadly agrees with expectations from detailed calculations and simulations (Walker *et al*., 2014, 2015).

### Gamma-ray Image Data processing

The quantification measurements for regions of interest (ROIs) provide a count of average gamma-rays detected s^−1^ 1280 mm^−2^ (these units are used since both ROIs are of that area on the face of the detector). High levels of gamma-rays were detected during the early time points, especially in *A. halleri*, and the gamma ray levels dissipated to a local minima around 3 h. In order to compare the dynamics of the Zn transport from root to shoot across the samples, the shoot ROI measurements were normalized to the local minima by subtraction. To calculate the rate of transport of Zn into the shoot ROI, the initial slopes of gamma ray build up for shoot ROI were calculated with the time points between 3 h and 24 h. To carry out a comparison of time points where the first differences in Zn accumulation into shoot could be observed, the data needed to be processed further. The imaging setup collects 8 × 10^6^ gamma-rays (over a typical time period of 1800 s) which summed together then constitutes a time point. Due to the difference in rates for each plant imaged, the time points for imaging are not matched sample to sample. In order to compare specific time points between the samples, values for 3 h, 4 h, 5 h, and so on until 12 h, were calculated using the two adjacent time points. This was done by calculating the slope and intercept between each pair of time points. Zn signal for each matched time point (*x*) was calculated using the formula *y = mx + b* (where *m* = slope and *b* = intercept from the values of two adjacent time points). Statistical analyses and time course plots were done in R version 3.4.2 (R Core Team, 2017) using agricolae (de Mendiburu, 2017), lsmeans (Lenth, 2016) and ggplot2 (Wickham, 2016) packages and wesanderson palette (Ram and Wickham, 2015).

## RESULTS

### Dynamics of Zn^2+^ movement in A. halleri vs. A. thaliana

The dynamics of Zn uptake and movement after Zn deprivation were visualized *in vivo* in intact plants of *Arabidopsis halleri* wild type (accession Langelsheim), the *A. halleri AhHMA4-*RNAi line 4.2.1 (in the Langelsheim accession background), and *A. thaliana* Col-0. First, the two species were grown to mature vegetative stage. *A. halleri* cuttings were grown hydroponically for 14 days to allow them to root. *A. thaliana* was grown on soil for 18 days, rinsed and transferred to Zn deprivation media (Hoagland media with no ZnSO4) for hydroponic growth. The plants were allowed to acclimate to Zn deprivation for 19–21 days. ICP-AES analysis of the Zn deprivation media showed that Zn concentration of the media was 0.12 µM. Immediately prior to imaging, the plants were transferred to resupply media with 1 µM ZnSO_4_ (normal concentration of ZnSO_4_ in Hoagland media and control concentration in (Talke *et al*., 2006)) and with 5.6 - 7.2 nM radioactive ^65^Zn. The plants were on the radiolabelled resupply media for 60 min, after which they were rinsed three times with non-radiolabelled resupply media before placing them on fresh non-radiolabelled resupply media for imaging with the UCD-SPI system. The time at which a plant was placed in the imaging system is considered the 0 h time point (Figures 1 and 2). Zn signal was measured in both the root region of interest (“root ROI”, blue box, Figure 1) and in the shoot region of interest (“shoot ROI”, red box, Figure 1). In *A. halleri* wild-type plants, the signal was detected in the root at 0 h, and then gradually moved up towards the shoot visibly at 12 h and 24 h, and subsequently appeared stationary in the upper part of the root from 36 h to 60 h (Figure 1A). In *A. halleri HMA4*-RNAi plants, the total gamma-ray signal never moved upwards from the root ROI, and instead moved slightly downwards in the root ROI over time, and the signal weakened considerably between 0 h and 12 h (Figure 1B). In *A. thaliana* wild-type plants, similarly to *A. halleri HMA4-RNAi* plants, no root-to-shoot movement of gamma-ray signal was visible, and the signal dissipated along the time course (Figure 1C). The radioactive signal in both the shoot and the root ROI started at their maximum for all *A. halleri* samples (Figure 2A, Supplementary Figure 1), and reduced to a local minimum at approximately 3 h. This effect is likely attributable to loosely bound ^65^Zn in the root apoplast, which is gradually desorbed into the external solution, thereby leaving the center of the field of view and reducing the likelihood of an emitted gamma ray being detected in the imaging system. In order to quantify and investigate root-to-shoot Zn transport dynamics, the focus was on the timepoints after the local minima, thus measurements were normalized to the time point at which the detected signal was the lowest close to 3 h. Shoot data corrected for the local minima is shown in Figure 2B.

**Figure 1.**
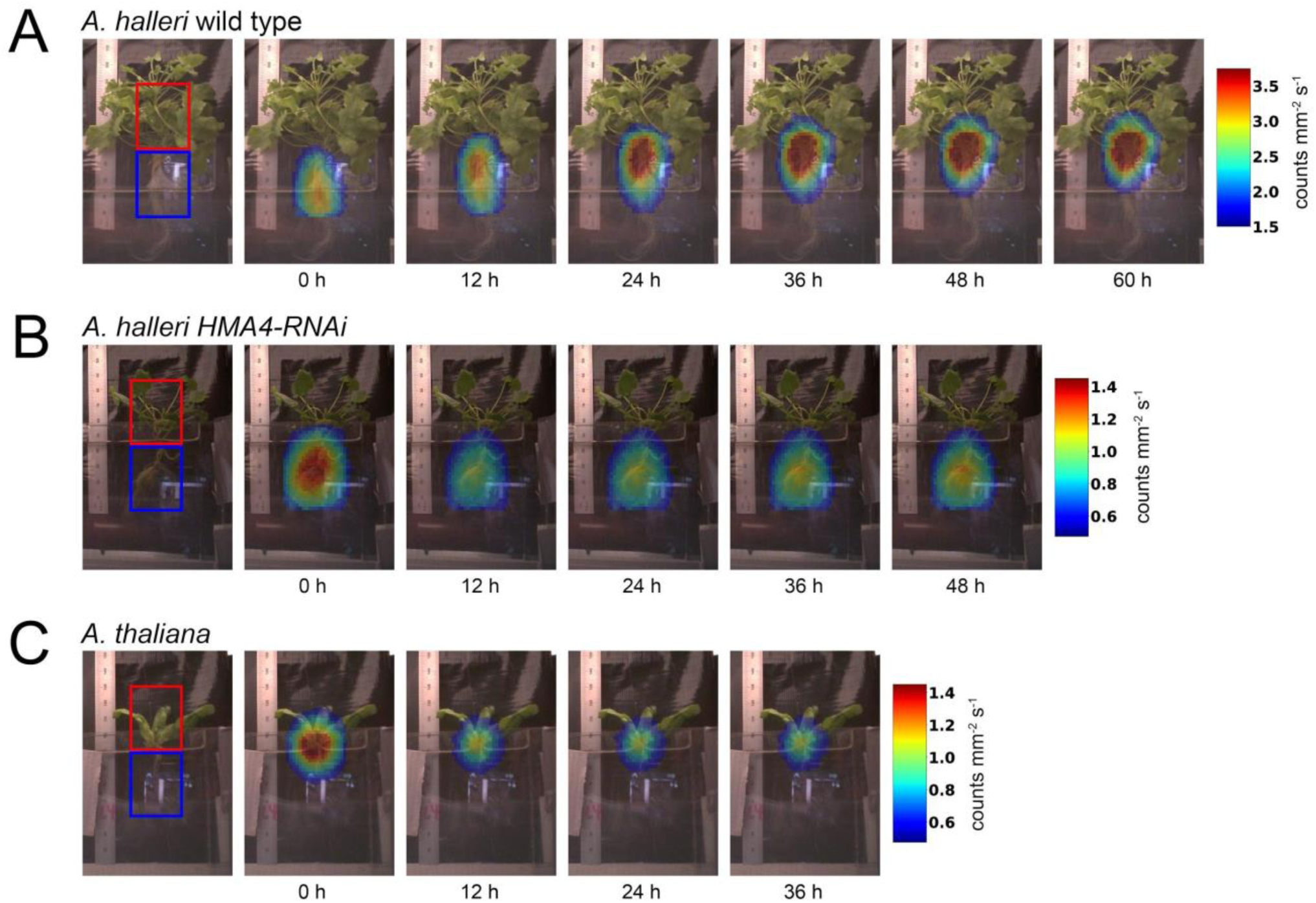
Visualization of whole-plant Zn resupply dynamics as detected by UCD-SPI. Zn dynamics were measured for *A. halleri* wild type (**A**), *A. halleri HMA4-*RNAi (**B**) and *A. thaliana* Col-0 (**C**) plants upon Zn resupply (1 µM ZnSO_4_) after 3 week Zn deprivation. Processed gamma ray detection data is overlaid with static pictures of the plants from which the data were collected. The red square represents the shoot ROI, and the blue square represents the root ROI.

At the time points after 3 h of re-supply of radio-labeled Zn, a continuous linear relative increase in the amount of ^65^Zn was observed in the shoot of *A. halleri* wild-type plants (Figure 2C). By contrast, in both *A. halleri HMA4-RNAi* and *A. thaliana* shoots, no additional ^65^Zn accumulated over the time period examined. On the contrary, the *A. halleri HMA4-RNAi* shoots lost further ^65^Zn signal after the initial local minimum, and from 7 h onwards this was statistically significant (Tukey’s HSD test, *p* < 0.05). The differential net ability of the three genotypes to take up Zn from the growth medium was determined through independent Zn resupply experiments. These experiments were conducted over the identical time period (24 h) to the gamma ray imaging experiment, but ICP-AES was used to measure the amount of Zn remaining in the growth media (Figure 2D). These data demonstrated that also at the whole-plant level, *A. halleri* wild-type plants took up significantly more Zn from the growth media than *A. halleri HMA4-*RNAi and *A. thaliana* (Figure 2D).

**Figure 2.**
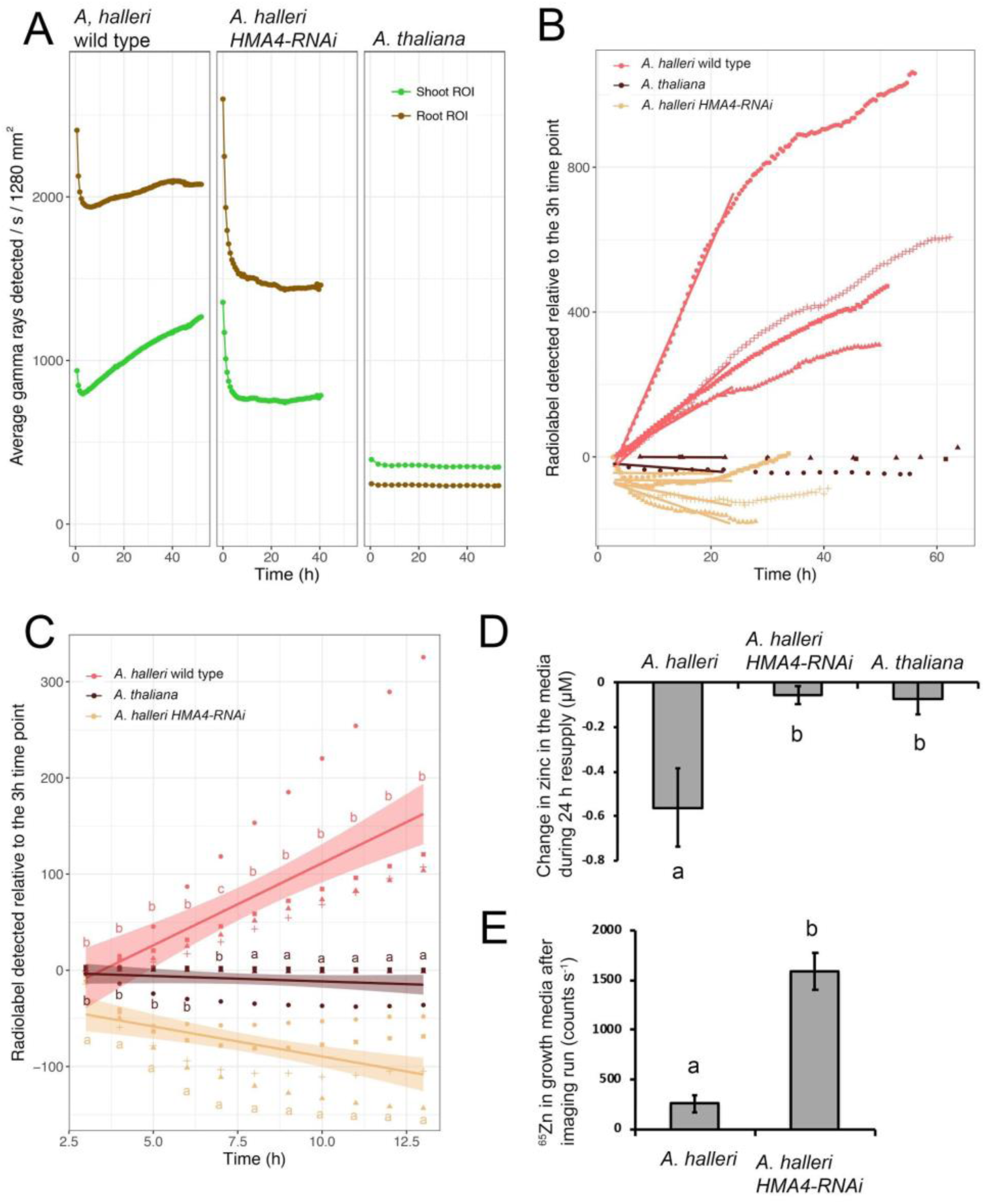
Quantification of Zn root-to-shoot transport dynamics. **A:** Examples of raw data points for shoot (green) and root (brown) ROI measurements across the imaging time lapse (average gamma-rays detected 1280 mm^−2^ s^−1^). For *A. halleri* wild type and the *A. halleri HMA4-*RNAi line, the 0 h time point shows a high level of gamma-ray detection that dissipates to a local minimum at 3 h. **B**: Gamma-rays in the shoot normalized to the 3 h local minima of the shoot ROI. In order to compare the dynamics of the ^65^Zn transport to the shoot across the samples, the values were normalized to the local minima. The slopes represent the rate of signal intensity change over time transport from 3 h to 24 h. The initial slopes are significantly steeper in *A. halleri* wild type compared to the two other genotypes (ANOVA, TukeyHSD, *p* = 0.0126). Pink data points: *A. halleri* wild type. Beige data points: *A. halleri HMA4-RNAi*. Brown data points: *A. thaliana*. Different symbol shapes indicate replicates (*n* = 4 for *A. halleri* wild type and *A. halleri HMA4-RNAi, n* = 3 for *A. thaliana*). **C**: Gamma-rays visualized in shoots as in (B), displayed at discrete matched time points. The gamma-ray imaging setup collects 8×10^6^ gamma-rays which then constitute a time point. Due to this, the time points for imaging are not matched sample to sample. In order to compare specific time points between the samples, values for 3 h, 4 h, 5 h, and so on until 12 h, were calculated using the two adjacent time points. The letters indicate samples that are significantly different between genotypes at each time point (*p* < 0.05, least-squares means). Lines represent the means and smoothed areas standard errors for each genotype. **D:** Analytic ICP-AES quantification of Zn in resupply media after 24 h resupply compared to 0 h. **E:** Gamma-rays detected in the imaging media after plant imaging (average gamma-rays detected s^−1^ 1280 mm^−2^). As there is no radiolabelled ^65^Zn in the imaging media prior to when the plant is placed for imaging, this represents radiolabelled ^65^Zn leached from the plant into the media. Letters (in panels D, E) indicate significant differences (*p* < 0.05, Student’s t-test) and error bars (panels D, E) indicate standard error.

The root ROIs were also analysed for change in ^65^Zn levels (Supplementary Figure S2). All three genotypes had identically no change in the ^65^Zn signal across the analysed time course (Tukey’s HSD test).

### Silencing of HMA4 completely abolishes root-to-shoot transport of Zn and results in Zn extrusion

The HMA4 transporter pumps Zn^2+^ from the root symplasm into the apoplastic xylem sap of *A. thaliana* (Verret *et al*., 2004). Strongly elevated expression of *A. halleri HMA4* was suggested to be responsible for the increased in root-to-shoot translocation of Zn in *A. halleri* relative to *A. thaliana (Hanikenne *et al*., 2008)*. This conclusion was drawn based on the quantification of shoot Zn concentrations after long-term growth in *HMA4*-RNAi lines and wild-type *A. halleri* and in *A. thaliana* Col-0 (Hanikenne *et al*., 2008). In the same experiment, root Zn concentration was elevated in some *A. halleri HMA4* RNAi lines relative to *A. halleri* wild-type plants and even relative to *A. thaliana (Hanikenne *et al*., 2008)*. In our whole-plant imaging system, the *A. halleri HMA4* RNAi line, shoot ^65^Zn levels in fact decreased over a 24 h time period (Figure 2B-C, Tukey’s HSD test, *p* < 0.05). Finally, the amount of ^65^Zn signal remaining in the growth media after the imaging run is six-fold higher after imaging of *A. halleri HMA4-*RNAi compared to wild-type *A. halleri* (Figure 2E), indicating a higher net loss of ^65^Zn from the *A. halleri HMA4* RNAi root including the apoplast into the surrounding media. This ^65^Zn represents Zn that was transferred into the imaging setup with the plant after the ^65^Zn treatment and root rinsing.

## DISCUSSION

### Advantages of gamma-ray imaging in intact plants

Aspects of metal uptake and homeostasis in plants may be understood well at the molecular level, but understanding of the whole-plant dynamics has lagged behind due to the limitations of traditional experimental approaches and imaging systems. Radiolabeled molecules are widely used to measure the transport dynamics in biological systems. In experiments with whole plants and radiolabeled molecules, the biodistribution of the radiolabel is most typically analyzed by plant dissection and counting in a well counter, and for β-emitters, by ashing the plant biomass and counting the radiolabel with a liquid scintillation counter. Several whole plant positron emission tomography (PET) imaging systems have been developed using ^11^C (Jahnke *et al*., 2009; Weisenberger *et al*., 2009; Kawachi *et al*., 2011), other groups have developed large scanners for β-imaging using ^32^P (Kanno *et al*., 2007), and autoradiography has been used to image radioisotope distribution of whole plants (Page and Feller, 2005). A scanner (Kawachi *et al*., 2011) has been used to perform PET imaging of ^65^Zn and ^107^Cd in rice plants (Fontanili *et al*., 2016; Suzui *et al*., 2017). Each of these nuclear imaging methods has drawbacks: for β-imaging, the range of β-particles can be too short to escape the plant; for PET, radioisotope lifetimes are short and the range of the positron can be too long for a thin, low density plant, limiting the yield of annihilation gamma-rays; and autoradiography is invasive and can require exposure times of weeks or months. Therefore, the UCD-SPI system used for nuclear imaging of gamma-ray emitters in plants has significant opportunity to contribute in a new way to transport studies. In particular, the uniquely high sensitivity of the system means that very small amounts (nCi) of radioisotopes may be imaged and followed over time. The system used here had the advantage that one entire time course can be recorded on a single plant. In time courses involving destructive sampling or imaging, data for the different time points are from distinct plant individuals, respectively, which generates substantial noise. This is particularly disruptive in experiments with non-model plants such as *A. halleri*, which show substantially larger variation in plant architecture between independently grown plants even of an identical genotype.

Common PET isotopes of interest for plant studies include several found in organic compounds: ^11^C (T_1/2_=20 min), ^13^N (T_1/2_=10 min), and ^18^F (T_1/2_=109 min). These atoms can be substituted into amino acids or sugars in plants to follow the natural *in situ* processes (Cherry *et al*., 2003). However due to the short lifetimes of PET radioisotopes, only short biological processes, such as photosynthesis, may be imaged. In contrast, single gamma-ray emitting radiotracers used in single photon emission computed tomography (SPECT) are typically metals and do not easily label organic molecules. However, many trace element metals are essential to a plant’s survival (Williams *et al*., 2000; Clemens, 2001). An active area of plant research is studying hyperaccumulation of metals in plants using a radioisotope of that metal; commonly studied metals include: Cd, Zn, Mn, Co, and Ni (Page and Feller, 2005; Krämer, 2010). Other potential applications for SPECT imaging include: studying plant ion transport in xylem (Macklon, 1970; Ueno *et al*., 2008), studying metabolic processes such as tracers for phloem transport (Omid *et al*., 2008), and studying signaling by labeled exogenous peptides or proteins (Santner and Estelle, 2009). One further advantage of imaging systems based on gamma-ray detection is the possibility of detecting the interactions of multiple radioisotopes simultaneously as the gamma-rays that they emit have distinct energies that can be distinguished from each other by the detector. For example, simultaneous imaging of ^65^Zn (gamma-ray 1116 keV) and ^109^Cd (22 keV) would enable teasing apart the competition dynamics in their uptake.

### Zn transport dynamics in A. halleri and A. thaliana

Zn uptake into the symplast in the outer root layers and loading into the apoplastic xylem stream are well understood on molecular level (Moreira *et al*., 2018). However, the dynamics of symplastic movement and patterning of the radial transport have thus far only been modeled to elucidate the timescales of these events (Claus *et al*., 2013). After xylem loading, the mass flow-mediated movement of Zn into the shoot inside the xylem is expected to occur within 30 min in *Arabidopsis*, as previously shown for water (Park *et al*., 2014) and Cd in xylem sap (Ueno *et al*., 2008). From previous SPECT imaging, we have shown that a pulse of radiolabelled pertechnetate (^99m^TcO^4-^) moving in the xylem stream reaches the shoot apical meristem of a 2 week old sunflower already in 5 min (Walker *et al*., 2015).

The rate-limiting step for root-to-shoot translocation of Zn was proposed to be xylem loading involving HMA4 transporters in both *A. halleri* and *A. thaliana* (Sinclair *et al*., 2007; Hanikenne *et al*., 2008). The dynamics of root-to-shoot Zn flux, however, have so far remained unclear in different species and transgenic lines. Estimates of Zn translocation rates from root to shoot were first obtained by spectroscopy methods of ashed shoot tissues. Early work with metal hyperaccumulator *Noccaea caerulescens* suggested that the speed of root-to-shoot Zn transport was between 20 and 60 h (Lasat *et al*., 1996; Lombi *et al*., 2001). Recently, positron imaging of Zn uptake estimated the time for Zn root-to-panicle transport in dwarfed mature rice to be 5.3 h (Suzui *et al*., 2017). Here, we have produced the first Zn root-to-shoot imaging data for *A. halleri* using UCD-SPI. Zn accumulates within the shoot of *A. halleri*, consistent with its ability to hyperaccumulate Zn, different from *A. thaliana*. The speed of Zn transport into the shoot in our data as observed with the smoothed standard error as well as differences with *A. thaliana* show clear shoot accumulation within 5–7 h, respectively (Figure 2C). These results are in line with previous reports for rice (Suzui *et al*., 2017). This contrasts strongly with the faster speed of the other xylem-transported compounds, such as water in *A. thaliana* (Park *et al*., 2014), Cd in *A. halleri (Ueno *et al*., 2008)* and pertechnetate in sunflower (Walker *et al*., 2015), all measured to reach the shoot in 30 min. It should be noted, however, that the experiments demonstrating water transport (Park *et al*., 2014) and Cd transport (Ueno *et al*., 2008) were carried out using decapitated stems and are thus destructive in nature, but also far more sensitive to small quantities than the method used here. The slower speed of Zn transport indicates that Zn loading into the xylem by HMA4 is slow and under tight control even in the metal hyperaccumulator *A. halleri*. Modelling the radial transport of Zn uptake has indeed indicated that HMA concentration is one of the key determinants of the uptake dynamics (Claus *et al*., 2013).

*HMA4* is critical to the ability of *A. halleri* to hyperaccumulate Zn. We tested the functional role of *HMA4* for *A. halleri* Zn translocation from root to shoot by imaging the Zn uptake dynamics of *A. halleri HMA4-*RNAi line relative to *A. halleri*. We found that the Zn signal in the shoot of *HMA4-*RNAi line did not increase over our 40-h imaging period, but conversely, we saw a continuous decrease in shoot Zn signal with significant differences observable at 3 h (Figure 2C). The lack of an increase in shoot Zn confirms that Zn loading into the xylem is abolished in the *HMA4-*RNAi plants (Hanikenne *et al*., 2008). The continuous decrease in the Zn signal in the shoot ROI seems to reflect bleeding of the strong early Zn signal from the root ROI into the shoot ROI. The dissipating signal through the *A. halleri HMA4-*RNAi time course could be due to apoplastic ^65^Zn adsorbed to the cell walls of outer root layers during the ^65^Zn pulse (Lasat *et al*., 1996) and not removed by the triple rinsing with Hoagland solution. This cell wall-adsorbed ^65^Zn would be desorbed into the growth medium during the imaging period by diffusion. The influx of Zn into the root symplasm is very tightly and rapidly regulated in Zn-concentration dependent fashion (Talke *et al*., 2006; van de Mortel *et al*., 2006; Claus *et al*., 2013). Without the loading of Zn into the xylem, Zn builds up in the root symplasm. In the case of *A. halleri HMA4-*RNAi, the symplasm could be saturated with Zn at 3 h after the resupply, leading to prevention of further uptake of the cell wall-adsorbed ^65^Zn and thus higher Zn desorption than Zn uptake into the symplasm.

Finally, we compared the dynamics of Zn movement in the Zn hyperaccumulator *A. halleri* with those in the related species *A. thaliana*, a non-metal hyperaccumulator. We found that Zn resupply after Zn deprivation in *A. thaliana* did not lead to detectable uptake or change of Zn in the shoot or the root ROI. Regardless of lack of hyperaccumulation, *A. thaliana* is still expected to uptake and transport Zn from 1 µM ZnSO_4_ resupply media. It is possible that the small size and flat rosette growth habit of *A. thaliana* affected our ability to detect Zn dynamics. Also, low abundance of HMA4 transporters in *A. thaliana* roots may lead to much slower dynamics that we were unable to capture.

The heavy metal imaging study presented here is of interest for phytoremediation applications (Robinson *et al*., 1998; Salt *et al*., 1998; Kärenlampi *et al*., 2000; Krämer, 2010; Sarma, 2011). Although most plants prevent the accumulation of heavy metals so as to avert toxicity, metal hyperaccumulators selectively extract high concentrations of metals from the soil into their shoots without incurring symptoms of toxicity (Baker and Brooks, 1989; Frérot *et al*., 2010). By using the heavy metal radiolabel ^65^Zn and the UCD-SPI imaging system, we gained a more detailed spatiotemporal understanding of the dynamics of metal movement into plants, which may be a path toward the use and understanding of metal hyperaccumulating plants for such advantageous applications.

## SUPPLEMENTARY DATA

Fig S1. Detected gamma-ray data for every replicate for both shoot and root ROI.

Fig S2. Gamma-rays detected normalized to the 3 h local minima in the root ROI.

## ACKNOWLEDGEMENTS

KK was supported by Finnish Cultural Foundation postdoctoral fellowship. KLW was supported by Office of Science (BER), U.S. Department of Energy. SMB was partially funded as an HHMI Faculty Scholar.

## SUPPLEMENTARY FIGURE LEGENDS

**Supplementary Figure S1.**
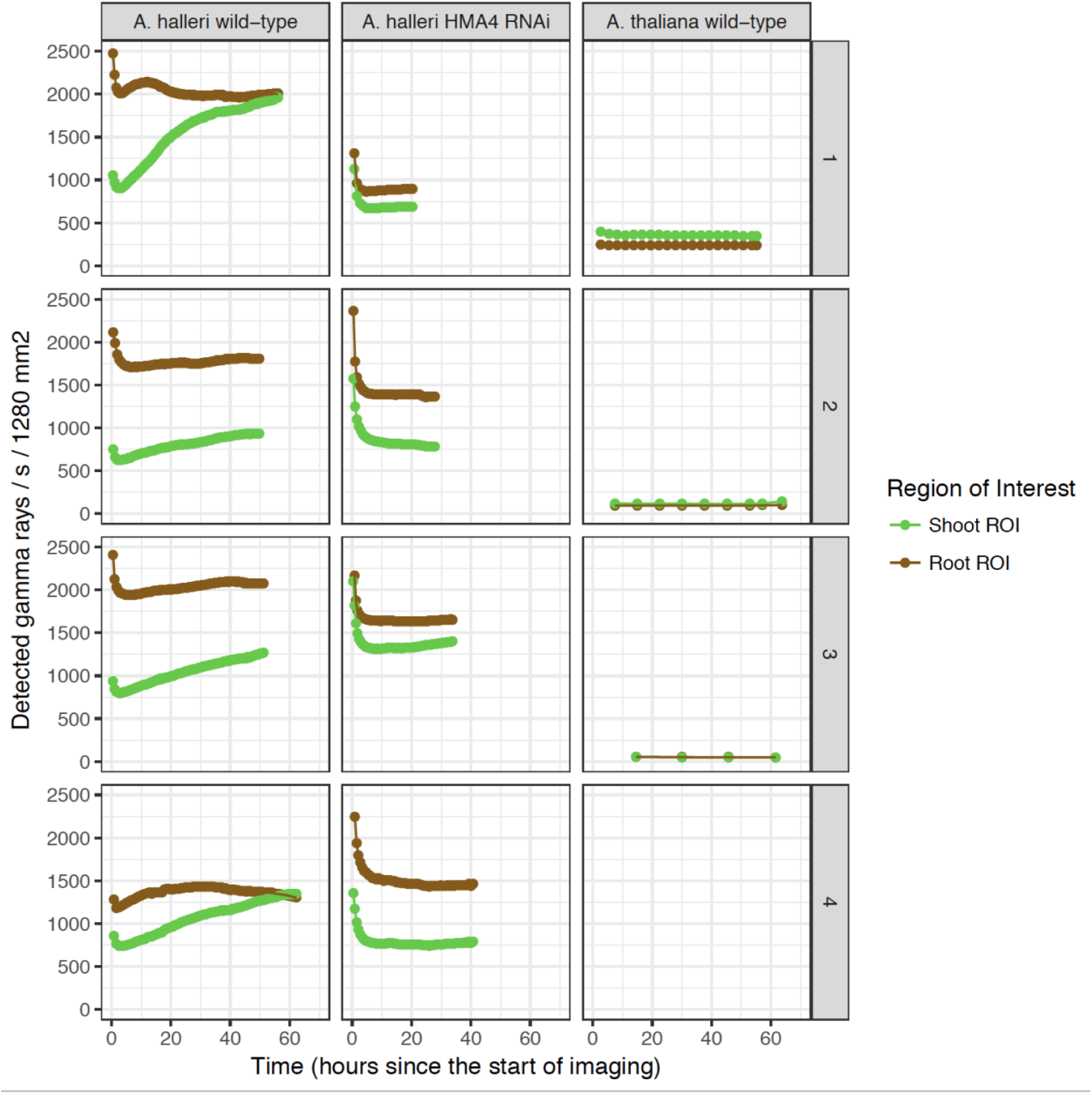
Detected gamma-rays from each genotype (columns) and replicate (rows) for root (brown) and shoot (green) ROIs.

**Supplementary Figure S2.**
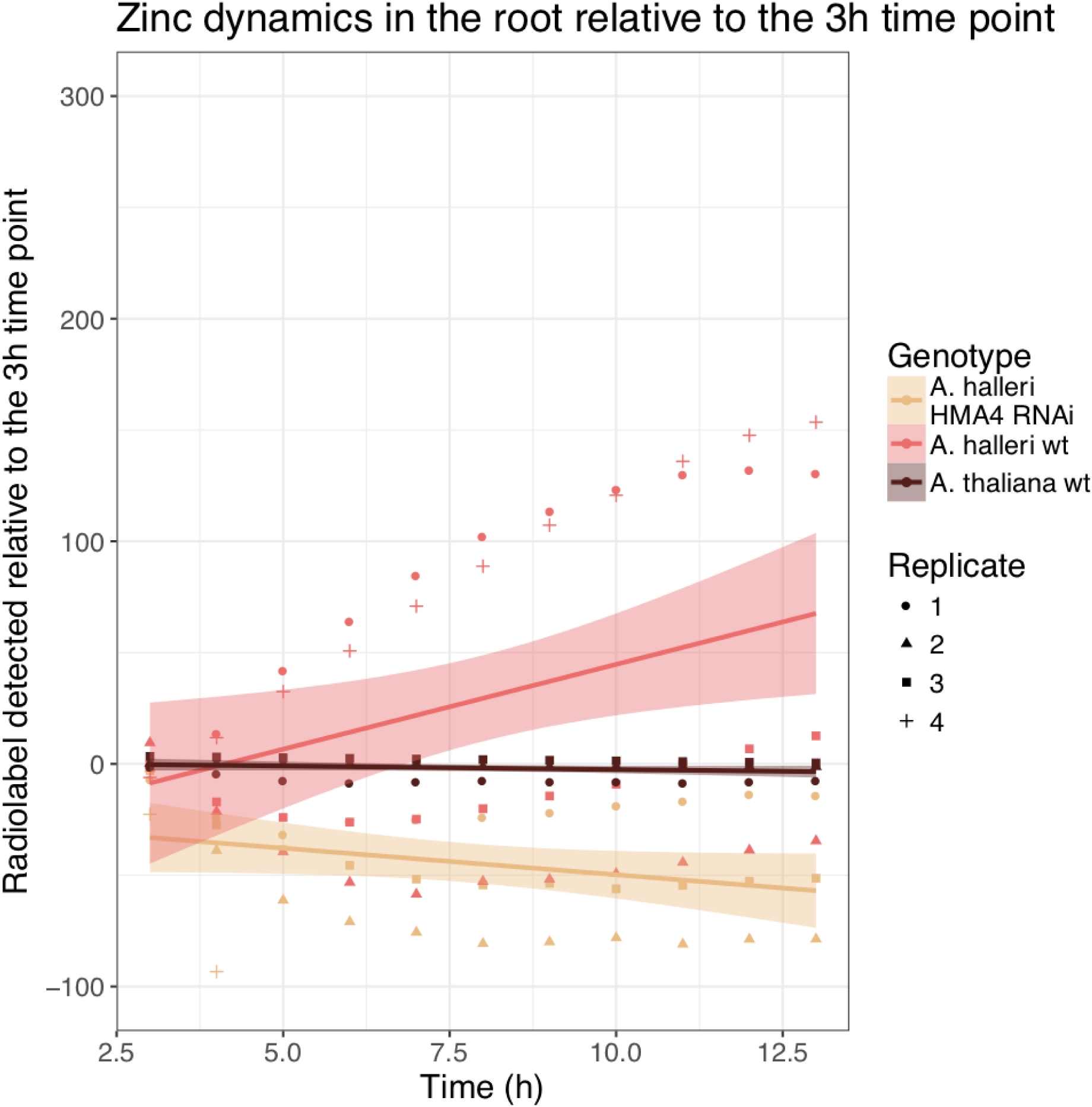
Gamma-rays detected normalized to the 3 h local minima in the root ROI. Pink data points: *A. halleri* wild type. Beige data points: *A. halleri HMA4-RNAi*. Brown data points: *A. thaliana*. Different symbol shapes indicate replicates (*n* = 4 for *A. halleri* wild type and *A. halleri HMA4-*RNAi*, n* = 3 for *A. thaliana*).

## Abbreviations

ICP-AES: = Inductively coupled plasma atomic emission spectroscopy
ROI: = region of interest
SPECT: = single photon emission computed tomography

